# Translate the untranslated: excising as a novel post-splicing event

**DOI:** 10.1101/2020.02.07.938738

**Authors:** Dmitry Y. Panteleev, Roman V. Reshetnikov, Nadezhda S. Samoylenkova, Nikolay A. Pustogarov, Galina V. Pavlova

**Author notes:** These authors contributed equally to this work (Dmitry Y. Panteleev, Roman V. Reshetnikov).

## Abstract

Maturing of a messenger RNA (mRNA) is a multi-way process producing mRNA variants through diverse splicing events, alternative polyadenylation, RNA editing, etc. Studying posttranscriptional processing of human cold-inducible RNA binding protein (CIRBP), we discovered yet another mechanism that could be added to this list. We named it excising, and it consists of low-accuracy post-splicing deletion of sequence regions of variable length. The main features of the excising process and putative members of corresponding multiprotein machinery were described with a series of cloning vectors and RNA-pulldown assay. Our results highlight a possible role of U-rich stretches and the proteins targeting such motifs in the discovered process. The discovered mechanism suggests the potential translation of 3’-untranslated regions, which may be an adjuvant way of CIRBP activity inhibition or generation of structural and functional diversity.

## Introduction

A mature eukaryotic messenger RNA (mRNA) starts its life-cycle as a primary transcript from genomic DNA consisting of coding (exons) and non-coding (introns) pieces. Besides that, mRNA is also framed with untranslated regions (UTRs) that contain various control elements regulating its translation. Further processing of a eukaryotic transcript includes 5’-capping, 3’-polyadenylation, and removing of introns (splicing). At the late stages of the lifecycle, mRNAs undergo degradation through deadenylation (1), nonsense-mediated decay (2), and other pathways.

There are several known mechanisms of proteome diversity generation in eukaryotic cells, which produce cell-, tissue- and condition-specific mRNA variants. The most common is a process known as alternative splicing, where exons are joined in different combinations forming alternative transcripts. Exon recognition depends on consensus sequences located at intron/exon junctions (3). Non-canonical splicing events are associated with the existence of similar motifs within introns as well as atypical splice sites, which generate cryptic exons (4), microexons (5), recursive splicing (6), circular RNAs (7), and other variations (8, 9). About 30 % of mRNAs contain alternative polyadenylation sites, which also contributes to transcripts variability (10). There is also a mechanism called RNA editing that involves incorporation of site-specific substitutions into RNA sequence, altering splicing patterns, and changing the coding sequence of mRNAs (11). Finally, a variety of protein products can be generated at the translation stage through the process known as recoding (12). It is associated with surrounding mRNA species that form competitive translation pathways.

Evidently, the eukaryotic apparatus for generating functional diversity is abundant. Nevertheless, when studying post-transcriptional processing of *CIRBP* gene in tumor cells, we encountered a process that does not fit into any of the previously described types. CIRBP is an 18-kDa protein that consists of an N-terminal RNA recognition motif (RRM) domain and a C-terminal arginine-glycine rich motif. CIRBP upregulation is observed in various tissues upon mild hypothermia, cold stress, UV radiation, mild hypoxia, and glucose deprivation (13). CIRBP serves its function by binding specific 51 nucleotides-long U-rich motifs in the 3’-untranslated region (3’-UTR) of mRNA transcripts (14, 15). This binding increases mRNA stability, consequently enhancing translation (14, 15). The role of CIRBP in tumorigenesis remains controversial. In some cases, it has an oncogenic function (16), while in other studies, it acts as a tumor suppressor (17).

According to our results, besides canonical alternative splicing, *CIRBP* has a splicing-independent low accuracy mechanism of mRNA diversity generation. We highlighted a possible role of U-rich stretches and identified the proteins putatively associated with this process. We believe that this previously undescribed mechanism is also inherent in other genes, which could give a fresh angle on eukaryotic transcriptome analysis. Surprisingly, we have also observed an increase in the transcript lengths of some synthetic constructs used in the study due to the insertion of coding sections of the same gene inside the maturing transcript. Here we describe the main features of these processes, which we named *excising* and *incising*, correspondingly.

## Materials and Methods

### A. Cell cultures, cloning, and sequencing

HEK293, MCF7, HEF, HCT116, B-16, RAG, and Sus/fp2 cell cultures were used in this study. The cells were grown in Dulbecco’s modified Eagle’s medium (PANECO, Russia) or Dulbecco’s modified Eagle’s medium/F-12 mix (PANECO, Russia) supplemented with 10% fetal bovine serum (Ther-moFisher, USA). The cells were passaged every week, at a split ratio of 1:2. Before lysis, cells were removed from the culture vial with the trypsin-EDTA solution (PANECO, Russia) and washed with PBS.

Total RNA was isolated using TRIzol reagent (ThermoFisher, USA) and treated with TURBO DNase (Invitrogen, USA) to remove genomic DNA according to the manufacturer’s protocol. Reverse transcription was performed with random hexamer primers using the MMLV RT kit (Evrogen, Russia). The obtained cDNA was used as a template in RT-PCR. RT-PCR was performed with Taq polymerase (Evrogen, Russia) or Phusion High-Fidelity DNA polymerase (ThermoFisher, USA) according to the manufacturer’s protocol. The primers used in this study are listed in Table 1.

**Table 1.**
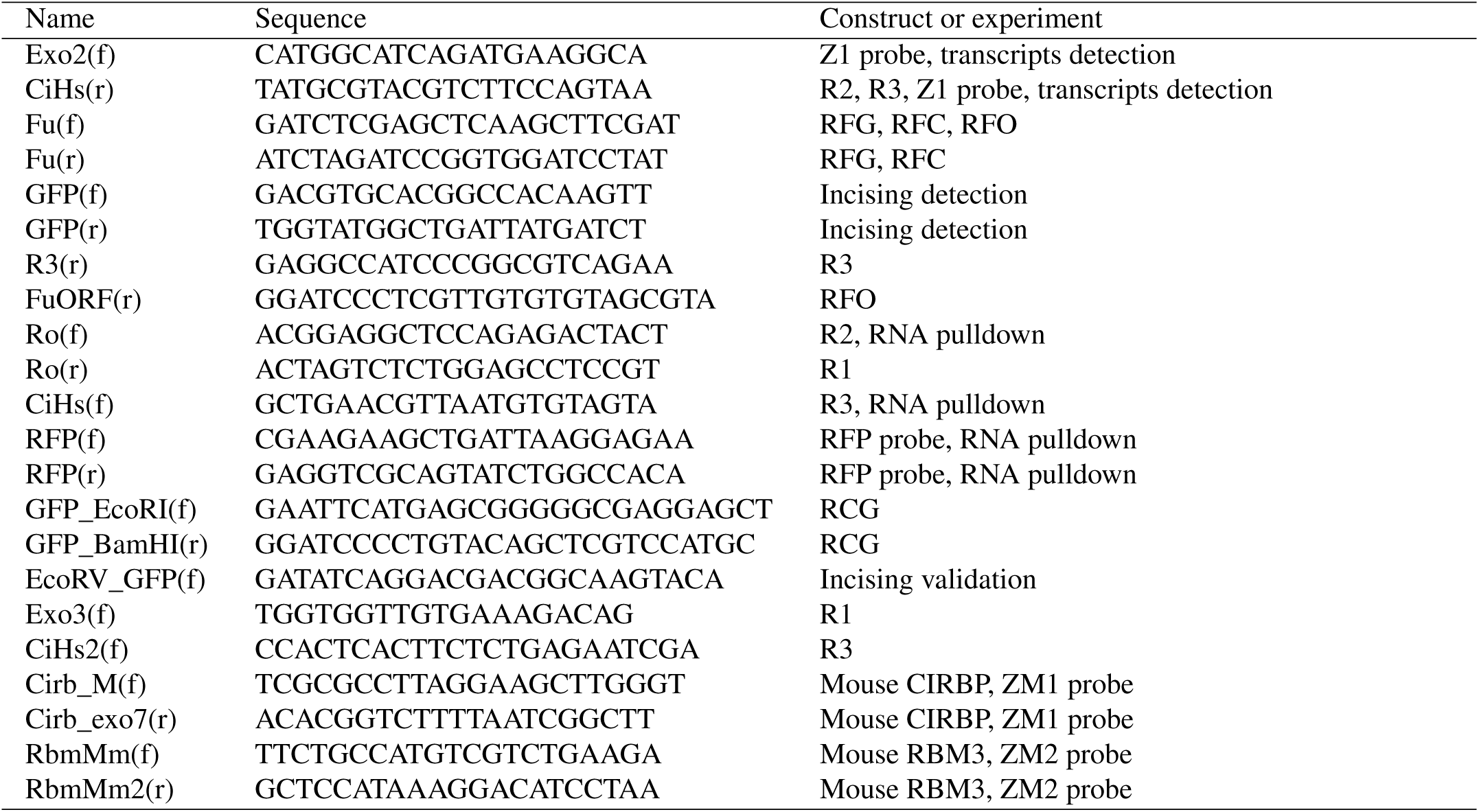
Primer sequences used in the study

For sequencing, PCR products were cloned into pGEM-T Easy Vector System (Invitrogen, USA). The sequencing was performed by standard Sanger sequencing at Evrogen, Russia facilities. Sequence alignment was performed with MAFFT (18) and visualized with AliView (19).

#### A.1. RFG, RFC, and RFO constructs

Sequences of *CIRBP* pre-mRNA, spliced mRNA (GenBank AN NM_001280.2), and spliced mRNA without UTR were cloned into HindIII and BamHI linearized pTagRFP-C vector using corresponding primers. For the RFG and RFC constructs, the primers set Fu(f) and Fu(r) was used. The RFO construct was created with a set of primers Fu(f) and FuORF(r). For the RF17 construct, the transcript sequence from clone 17 (Supplementary Note 1) was extracted from pGem vector with HindIII and BamHI restriction enzymes and inserted into pTagRFP-C vector using the same sites.

#### A.2. RCG construct

EcoRI restriction site was introduced at the position 1256 of the RFC construct with site-specific mutagenesis. *GFP* coding sequence was flanked with restriction sites for EcoRI at the 5’-end and BamHI at the 3’-end (after stop-codon), using PCR with a set of primers GFP_EcoRI(f) and GFP_BamHI(r). RCG vector was digested with EcoRI and BamHI, and the *GFP* sequence was inserted into it using the same sites to form the RCG construct.

#### A.3. R1-R3 constructs

EcoRV restriction site was introduced at the position 980 of vector pTagGFP2-N with site-specific mutagenesis. The vector was then digested with EcoRV, followed by T-tailing for AT-cloning of corresponding PCR products.

### B. Transfection

80% confluent monolayers were transfected using TurboFect transfection kit (Thermo Scientific, USA) according to the manufacturer’s protocol. The consequent manipulations were performed 24-48h after transfection. Phase-contrast and fluorescence microscopy were performed by CX41 microscope (Olympus, Japan).

### C. Southern blotting

PCR products were transferred to a positively charged nylon membrane (Roche, Switzerland) using capillary transfer under denaturing conditions. The probe was obtained with primers Exo2(f) and CiHs(r) and labeled using the Biotin DecaLabel DNA Labeling Kit (Fermentas, Lithuania). Hybridization was performed in the Church-Gilbert hybridization buffer.

### D. RNA-Seq data analysis

Illumina RNA-Seq paired-end reads obtained from HEK293 cells (ENA AN PR-JNA245463) were preprocessed using FASTX-Toolkit (20), and the longest *CIRBP* transcript was assembled with DART algorithm (21) using *CIRBP* genomic sequence (Entrez GeneID 1153) as a reference. Unipro UGENE tool (22) was used for visualization of the alignment.

### E. Northern blotting

Northern blotting was performed according to the standard protocol (23) using a positively charged nylon membrane (Roche, Switzerland). Before the transfer, a denaturing agarose gel electrophoresis was performed with 10 *µ*g of sample per lane loaded to gel. *RFP* RNA-probe was used for bands identification. The probe was labeled with biotin using T7 RNA Polymerase-Plus Enzyme Mix (Thermo Scientific, USA) by the addition of biotin-labeled UTP according to the manufacturer’s protocol.

### F. RNA-pulldown assay

The RNA-pulldown assay was performed as described elsewhere (24) with minor modifications. *CIRBP* sequence flanked by Ro(f) and CiHs(r) primers (or *RFP* sequence for the control system) with a 4-fold repeat of S1m aptamer (24) at the 3’-end was cloned into vector pGEM-T Easy Vector System (Invitrogen, USA). The linearized construct with blunted ends was used for in vitro transcription with T7 RNA Polymerase-Plus Enzyme Mix (Thermo Scientific, USA) according to the manufacturer’s protocol. Roughly 100 *µ*g of resulting RNA was immobilized on 50 *µ*l of High Capacity Streptavidin agarose (Thermo Scientific, USA). Cell lysate lysed from 0.4 g of HEK293 cells with lysis buffer (20 mM HEPES (pH 7.9), 150 mM NaCl, 5 mM MgCl2, 2mM DTT, 1% NP40, 5% glycerol, 1% PMSF, 1x Halt protease-inhibitor single-use cocktail (Thermo Scientific, USA) and 10 *µ*l RNase inhibitor Ribolock (Thermo Scientific, USA) for 15 min at 4 °C. After that, the sample was centrifuged for 10 min at 21000 g, and the supernatant was collected and added to the column. After 3 hours of incubation at room temperature with constant mixing, the column was washed three times with the lysis buffer, three times with high-salt buffer (20 mM HEPES (pH 7.9), 300 mM NaCl, 5mM MgCl2, 2mM DTT) and three times with low-salt buffer (20 mM HEPES (pH 7.9), 30 mM NaCl, 5mM MgCl2, 2mM DTT). After that, elution was performed with RNAze A (Thermo Scientific, USA), dissolved in the low-salt buffer to a concentration of 100 ng/*µ*l. The eluted fraction was precipitated with Ready Prep 2-D cleanup kit (Bio-Rad, USA). The precipitate was dissolved in O’Farrell lysis buffer and analyzed with 2D polyacrylamide gel electrophoresis. For identification of the RNA-binding proteins, the gel was stained using Coomassie Brilliant Blue G-250 (Helicon, Russia). The Coomassie-stained spots were then cut out from the gel and analyzed with MALDI-tof mass spectrometry. Functional enrichment analysis was performed using Gene Ontology (25, 26) and Pathway Commons (27) resources.

### G. 2D electrophoresis and western blotting

Protein lysates dissolved in O’Farrell buffer were analyzed by isoelectrofocusing, carried out in glass tubes filled with polyacrylamide gel using a mixture of ampholytes 3/10 and 5/8 (Bio-Rad, USA) overnight at 4 °C. PageRuler Plus Prestained Protein Ladder (Fermentas, USA) was used as a molecular weight marker. The second-dimension separation was performed in SDS 5-10% PAGE. For western blotting, the gel was transferred to a nitrocellulose membrane Hybond ECL (Amersham, USA).

After the transfer, the membrane was blocked for 1 hour at room temperature by x1 blocking reagent (Roche, Germany) dissolved in TBS-T buffer. After that, the membrane was incubated with primary antibodies overnight at 4 °C. As primary antibodies, we used rabbit polyclonal Anti-tRFP and Anti-TurboGFP(d) antibodies (Evrogen, Russia) diluted at 1:7000. As secondary antibodies, we used HRP-conjugated goat anti-rabbit antibody (Santa Cruz Biotechnology, USA) diluted at 1:10000. After washing, the results were detected with ECL Prime Western Blotting Detection Reagent (GE Healthcare, USA).

For the RCG construct experiment, pre-washed cells were lysed in x1 Laemmli buffer and analyzed with one-dimensional PAGE followed by western blotting using Anti-TurboGFP(d) antibodies (Evrogen, Russia).

## Results

### H. A variety of *CIRBP* isoforms in cancer and normal cells

To study the diversity of *CIRBP* transcript variants in cancer, we isolated total RNA from human glioma cell culture Sus/fP2 (28). After that, we used RT-PCR with a set of Exo2(f) and CiHs(r) primers specific to the start of *CIRBP* coding sequence and the end of 3’-UTR, correspondingly, to detect the transcripts. The resulting PCR products were blotted to a nylon membrane, and southern hybridization with the Z1 probe was performed (Figure 1a). We expected to observe PCR products of the known *CIRBP* transcript variants: Gen-Bank AN NR_023312.3, NR_023313.3, and NM_001280.2 with predicted lengths of 1531, 1424 and 1156 bp, correspondingly. Contrary to that, the probe exposed a great variety of detectable bands with electrophoretic mobility in the range of 200 bp to roughly 1200 bp (Figure 1a).

**Fig. 1.**
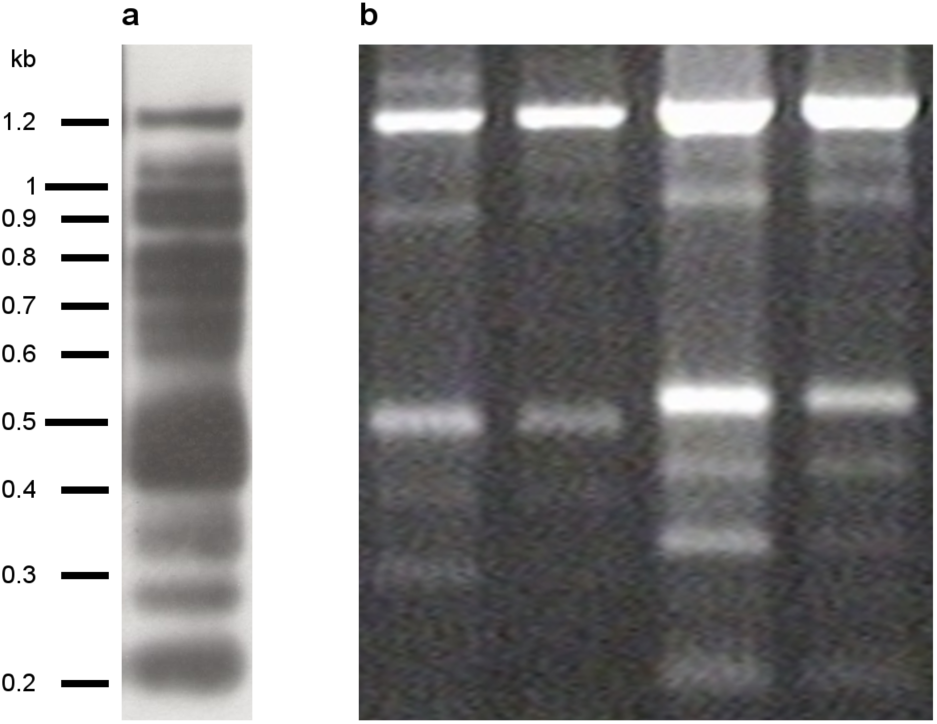
CIRBP transcripts diversity. **a**: southern blot analysis of RT-PCR products from Sus/fP2 cell culture; **b**: electrophoresis analysis of *CIRBP* transcripts in different cell lines. From left to right: MCF7, HCT116, HEK293, HEF. Note that the relative quantity of transcripts between individual bands may be affected by the PCR amplification effect.

To verify the southern blotting results, we performed the same analysis on cancer and normal cell lines MCF7 (breast cancer), HCT116 (colorectal cancer), HEK293 (embryonic kidney), and HEF (embryonic fibroblasts). In all the studied samples, we obtained the same pattern with varying intensity of individual bands (Figure 1b).

For further analysis, we selected two bands with electrophoretic mobility of 500 bp (500 series) and 1000 bp (1000 series) to explore sequence differences between the bands (Figure 1). The PCR products were cloned into a pGEM-T vector for a Sanger sequencing. Sequence analysis revealed that the variety of newly discovered isoforms corresponds to unconventional deletion variants of the known *CIRBP* gene transcripts (Figure 2a). In 500 series, the single continuous section of a transcript was excised (Figure 2a, b). However, some variants in the 1000 series contained several intraexonic deletions (Figure 2a). The full sequences are provided in Supplementary Note 1.

**Fig. 2.**
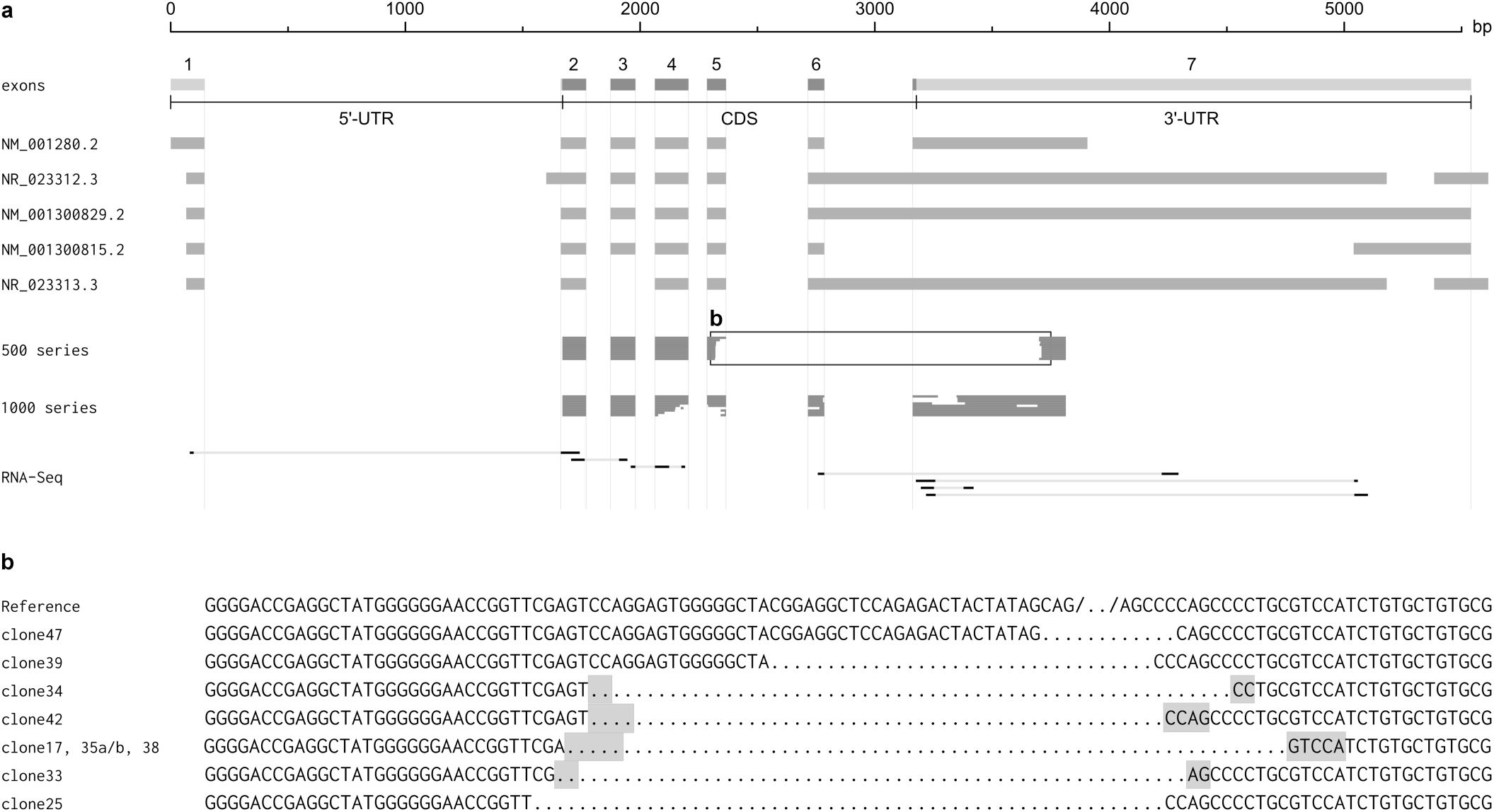
Alignment of CIRBP transcript variants. **a**: scheme of *CIRBP* transcript variant sequences. The list of RNA-Seq reads in order of appearance is provided in Supplementary Note 1. **b**: sequence alignment for the outlined region of 500 series from panel **a**. The sequence segments that may be aligned with the reference sequence both before and after an excised region are marked gray.

### I. Excising is a post-splicing event

The observed transcripts could be a result of either alternative splicing of pre-mRNA or some post-splicing event. To investigate the possibilities, we constructed fusion expression vectors containing a red fluorescent protein (RFP) sequence. The sequence of the first vector named RFG also included *CIRBP* pre-mRNA (Figure 3a), and the sequence of a mature transcript (Gen-Bank AN NM_001280.2) was cloned into the second construct named RFC (Figure 3a). The expression vectors were then transfected into HEK293 cells.

**Fig. 3.**
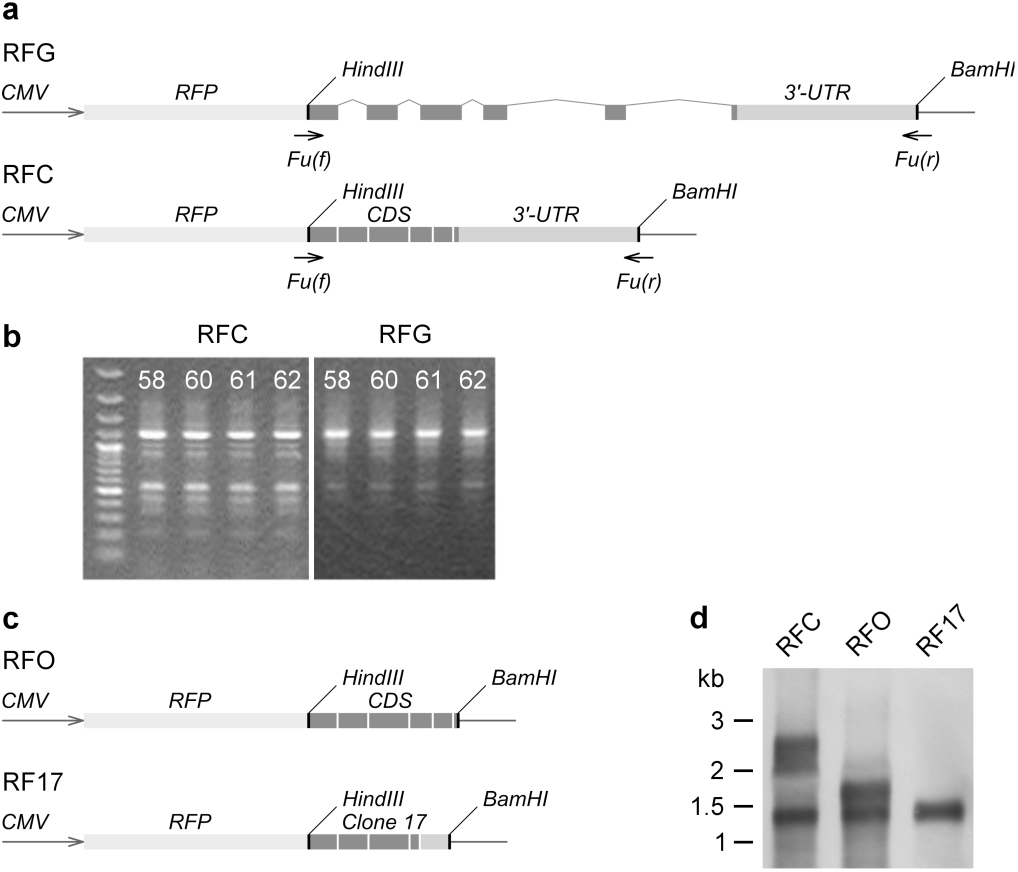
The place of excising in the flow of genetic information. **a**: schemes of RFG and RFC construct sequences. **b**: RT-PCR products of total RNA from RFG and RFC-transfected cells at different primer annealing temperatures, ^*°*^C. Molecular weight marker: GeneRuler 100 bp Plus DNA ladder (Thermo Scientific). **c**: schemes of RFO and RF17 expression vectors. **d**: northern blot analysis of HEK293 cells transfected by RFC, RFO, and RF17 constructs.

After 24 hours, we isolated total RNA from transfected cells and treated it with TURBO DNase (Invitrogen). To detect the transcript variants, RT-PCR was performed. We used a set of Fu(f) and Fu(r) primers targeting the sites starting 10 bp upstream or downstream of the 5’- and 3’-ends of *CIRBP* mRNA, correspondingly (Figure 3a). The patterns of the PCR products were identical between the RFG and RFC constructs (Figure 3b). Note that the patterns were also similar to the variety of bands observed for the native *CIRBP* gene transcripts (see Figure 1). According to these results, the transcript variants variety did not depend on the expression construct.

We also studied the putative effect of splicing on the observed phenomenon by repeating the experiment of HEK293 transfection with the RFG and RFC vectors in the presence of spliceosome inhibitor isoginkgetin (29). Isoginkgetin did not alter the southern blot patterns of the PCR products (Data not shown, but available under request). Together, these results may indicate that excising is a splicing-independent post-splicing event.

Sequencing of the PCR products revealed a low accuracy of the excising process; some of the sequences differed from each other by 4-7 nucleotides (see Figure 2, clones 25, 33, and 34). There were no consensus motifs at the junctions between the removed and retained sites.

The sequences also demonstrated that excising is not a recursive process of consecutive removal of sequence segments. The 500-series family of shorter sequences consisted of transcript variants containing intact exon 4 as well as a 5’-part of exon 5. The 1000-series collection of longer sequences had these regions heavily edited or removed (see Figure 2). This difference could mean that the observed pattern is the result of alternative processing of one or more initial sequences rather than a one-way process of stepwise shortening of a primary transcript. According to these results, excising is not a part of mRNA decay pathways.

We have also analyzed RNA-Seq data for HEK-293 to support our findings. According to the RNA-Seq data, excising is a relatively rare process. Out of 36204 paired-end reads mapped to the full-length sequence of *CIRBP* gene, only seven had characteristic features of excising (see Figure 2a). On the other hand, it is difficult to estimate the frequency of the excising process from a single RNA-Seq experiment. In our sequencing data excising occurred in exons 4, 5, 6, and 7. In the 500-series family of sequences, it targeted exons 5 and 7, with exon 6 removed. The seven reads from the RNA-Seq data showed that excising can occur in all *CIRBP* exons (Figure 2a). Five out of the seven reads resembled the excising pattern of the 1000-series sequences family, namely intra-exonic deletions inside exons 4 and 7. Note that none of the reads were compatible with the 500-series excising pattern, despite the relative abundance of this family according to RT-PCR results (see Figure 1).

### J. Excising occurs at the RNA level

To examine whether excising is an RT-associated process, we studied it on the RNA level using three fusion expression vectors: RFC, RFO, and RF17. All three vectors contained RFP at the 5’ end followed by different transcript variants of *CIRBP* gene. The RFO construct differed from RFC by the absence of 3’-UTR. The sequence of excised transcript variant from clone 17 (see Figure 2) was cloned into the vector RF17 instead of the conventional transcript (Figure 3c).

HEK293 cells were transfected with the expression vectors; after 24 hours, the polyA RNA was extracted from cell lysates and treated with TURBO DNase. The resulting RNAs were blotted to a nylon membrane, and northern hybridization with an *RFP* probe was performed. The probe exposed a band with a length of 1300 bp in all three samples (Figure 3d), while the lengths of conventional transcripts of RFC and RFO vectors were supposed to be about 2400 bp and 1600 bp, correspondingly. Note that the design of the RFO vector eliminates the possibility of a similar sequence for the band between the samples. It might mean that excising is initiated by an area that is not restricted to either coding sequence or 3’-UTR.

RFC-transfected cells, besides conventional and 1300 bp-long transcripts, contained an additional band with a length of about 2000 bp (Figure 3d). These results suggest that excising addresses RNA directly. Note that the difference in patterns between southern and northern blotting results (see Figure 1, 3d) is most likely related to PCR amplification of cDNAs, highlighting minor transcript fractions.

### K. A possible role of U-rich stretches in excising

We probed the boundaries of the excising mechanism by detection of the minimal sequence sufficient for the effect to occur. For that, we used a set of constructs based on the pTagGFP2- N expression vector. The set consisted of: (i) vector R1, containing 3’-end of *CIRBP* coding sequence and a major part of 3’-UTR; (ii) vector R2 with cloned segment of 3’-UTR, and (iii) vector R3, containing 5’-part of vector R2 (Figure 4a). HEK293 cells were transfected with the expression vectors, and after 24 hours of incubation, the total RNA was isolated and treated with TURBO DNase. We used RT-PCR with the corresponding sets of primers (see Table 1) to detect the transcripts. The resulting PCR products were blotted to a nylon membrane. Southern hybridization with the *GFP* probe revealed the excising-specific pattern for R1 and R2 vectors, but not for R3 (Figure 4b). R2 and R3 sequences differed in a section with a length of 199 nucleotides with a very high AU content (63.3 %). In terms of mono-, di- and tri-nucleotide frequencies, U nucleotides dominated the sequence of the section (45.2 %, 61.0 %, and 43 % for U, UU, and UUU, correspondingly). Note that there were no miRNA binding sites within neither R2 nor R3 sequences. U-rich stretches are known to provide structural destabilization of mRNA, being the targets of various proteins that can selectively bind these elements (31). It could be the basis of the excising mechanism.

**Fig. 4.**
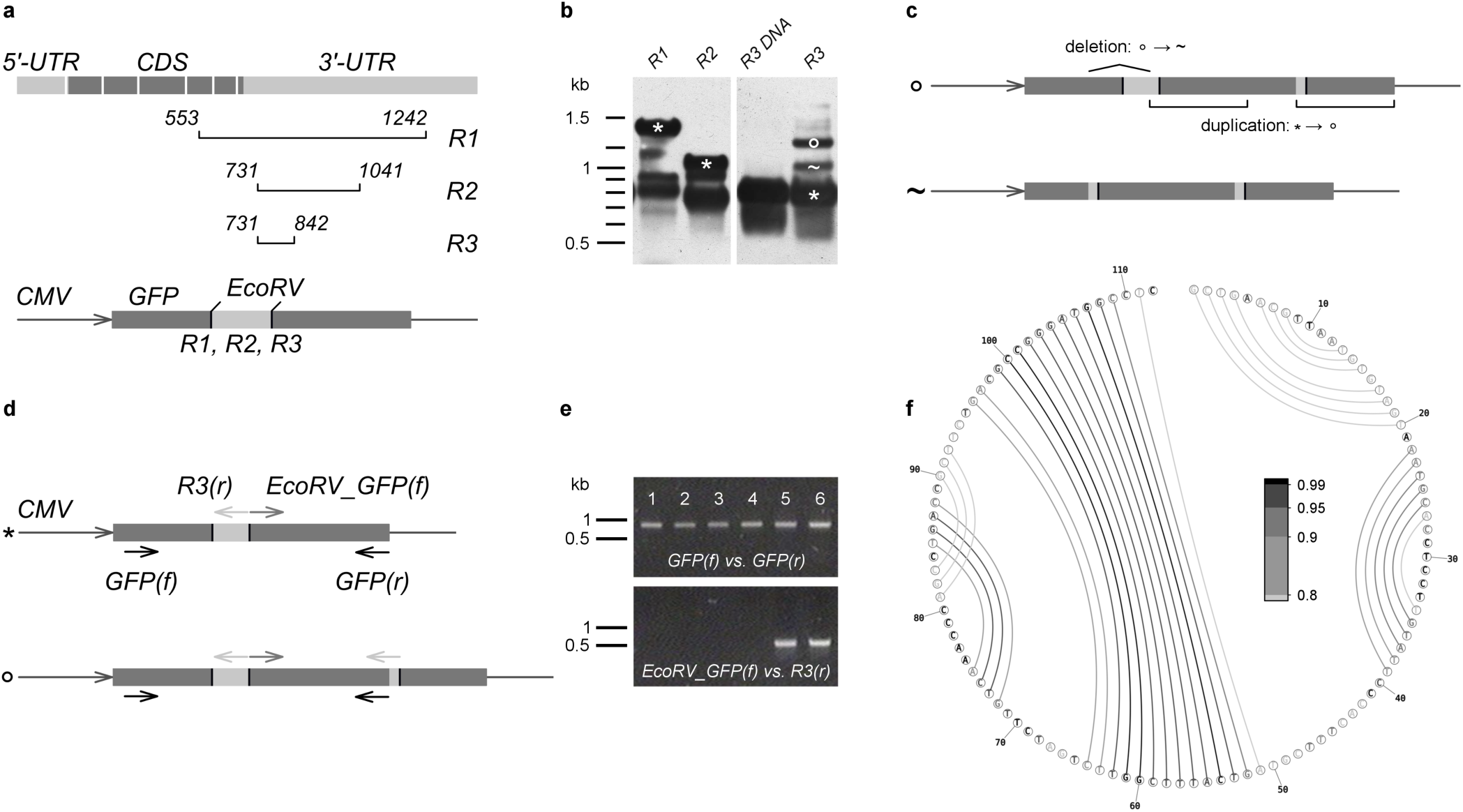
Chasing an excising-initiating sequence, incising detection and validation. **a**: design of R1-R3 constructs. Top: scheme of *CIRBP* transcript variant NM_001280.2 with marked regions used for vectors R1-R3 creation. For that, the marked regions were cloned into a pTagGFP2-N expression vector (bottom). **b**: results of southern hybridization of total RNA RT-PCR products after HEK293 transfection with vectors R1, R2, and R3 (lanes 1, 2, and 4, correspondingly). Lane 3: PCR product of R3-transfected HEK293 total DNA. Symbols “*” denote bands with expected lengths. **c**: sequence diagrams of bands “∘” and “∼” from the panel **b**. The full sequences are provided in Supplementary Note 2. **d**: design of incising validation experiments. Primers EcoRV_GFP(f) and R3(r) should result in a PCR product only in case of sequence duplication. **e**: PCR results for different matrices. Lanes 1, 2: initial R3 construct; lanes 3, 4: HEK293 genomic DNA extracted after transfection, lanes 5, 6: cDNA from HEK293 total RNA extracted after transfection. **f**: secondary structure prediction of R3 sequence according to ProbKnot utility of RNAstructure software package (30). The color-coding of base-pair probabilities is given in the center. All pairs with probabilities ≤ 0.8 are colored with light-gray.

### L. Incising

The surprising observation was the appearance of longer than expected PCR products in R3-transfected cells (Figure 4b). Note that such bands were not unique for the R3 vector, but the absence of excising in this system made them more prominent. According to sequence analysis of the additional bands, the longest amplicon (Figure 4b, variant “∘”) was a product of duplication of a 305 bp region containing 30 nucleotides from the 3’-part of *CIRBP* sequence and an adjacent segment of GFP sequence with separating them 3’-part of EcoRV restriction site (Figure 4c). The second in length amplicon (Figure 4b, variant “∼”) was a deletion product of the variant “∘”. Here, the upstream adjoining region of the duplicated site with the length of 106 bp was removed from the sequence (Figure 4c, variant “∼”).

We explored at what stage the sequence variation takes place using a set of inverted primers EcoRV_GFP(f) and R3(r). For the conventional sequence of vector R3, the set should not result in a PCR product. In the case of the duplication (variants “∘” or “∼”), we expected to observe a generation of a 500 bp PCR product (Figure 4d). As a control, we used a pair of primers GFP(f) and GFP(r) with a 900 bp PCR product (Figure 4d). We explored the results of PCR amplification at three stages of the experiment. As a DNA matrix, we used (i) the initial R3 construct; (ii) DNA extracted from cells after transfection, and (iii) cDNA from total RNA isolated from cells after transfection. All the matrices provided PCR products of the expected length for the control pair of primers (Figure 4e, top), whereas the inverted pair worked only on the third system (Figure 4e, bottom). The results indicate that the duplication occurs either at the RNA level or during RT. We named this process incising and plan to explore this phenomenon in future studies.

### M. Excising results in protein products

To examine the possibility of mRNAs translation after excising, we used two fusion constructs, RFC and RFO. We believed that in the case of excising, there would be a difference in protein products’ amount and diversity between the two constructs.

We performed a 2D western blot analysis of transfected cells proteome using RFP specific antibodies for the identification of expressed proteins (Figure 5a, b). The expression of RFP-labeled products in the RFO-transfected cells was substantially lower. Still, there was a difference in the protein composition judging by the protein spots (Figure 5a, b, spot 25). To detect the minor protein fractions in the RFO-transfected system, we used longer exposure time. It revealed additional protein spots uncharacteristic of RFC-transfected cells (Figure 5c, spots 20-26).

**Fig. 5.**
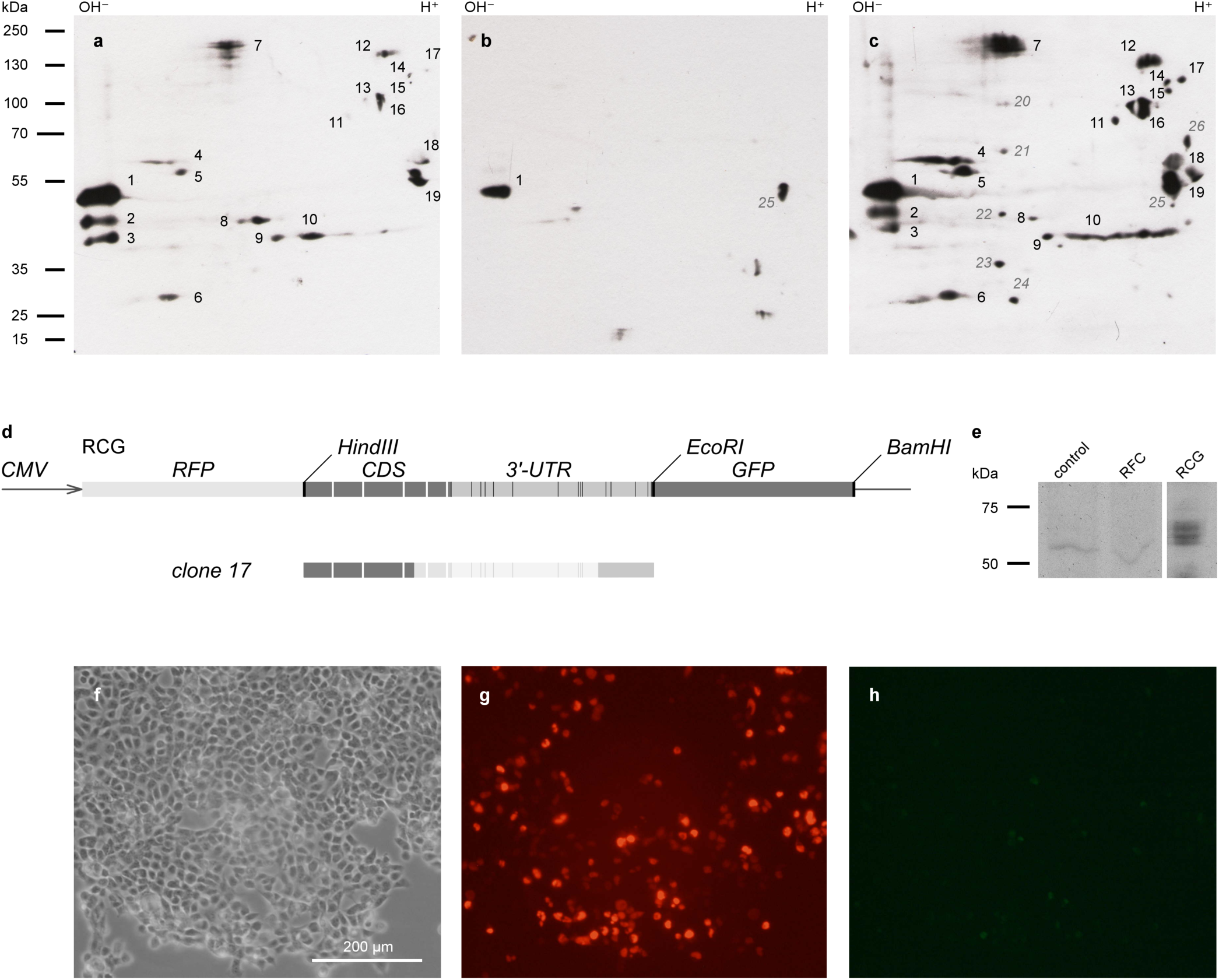
Translation of mRNAs after excising. **a-c**: western blot analysis of HEK293 cells transfected by RFC (panel **a**) and RFO (panels **b, c**) constructs. The numbers denote individual spots. The numbers for spots that are different between the two systems are in italic and colored gray. **d**: schemes of RCG construct (top) and transcript variant from clone 17 (bottom) sequences. The excised region in the latter is thinned down. Stop-codons are shown with thin black bars. **e**: western blot analysis of HEK293 cells lysates with GFP-specific antibodies. As a control, we used total lysate of non-transfected cells. **f-h**: imaging of RCG-transfected cells with phase contrast (panel **f**), RFP fluorescence (panel **g**), and GFP fluorescence (panel **h**).

To further validate the post-excising translation of mRNAs, we constructed a fusion expression vector RCG containing *CIRBP* transcript sequence flanked by RFP- and GFP-coding sequences at the 5’- and 3’-ends, correspondingly (Figure 5d, top). There were 15 stop codon sites in the same open reading frame (ORF) between the CIRBP- and GFP-coding sequences that prevented translation of the latter. However, we observed the deletion of the stop codon-containing sequence segments in the transcript variants affected by excising (Figure 5d, bottom) and ORF shifts (Supplementary Note 1), which could lead to the expression of GFP.

The RCG expression vector was transfected into HEK293 cells. There was a possibility of an IRES-like motif occurrence triggering the *GFP* translation instead of excising. We examined the possibility by western blotting of the HEK293 total lysate using GFP specific antibodies (Figure 5e). The observed bands were consistent with predicted electrophoretic mobilities of RCG protein products after excising. We also analyzed RCG protein products with phase-contrast and fluorescence microscopy (Figure 5f-h). We used RFP fluorescence as a control of transfection efficiency (Figure 5g). About 10-20% of transfected cells also exhibited a green fluorescence (Figure 5h), which indicates both the fact of *CIRBP* sequence excising and GFP expression in these cells. An intriguing consequence of this experiment is that excising allows translation of 3’-untranslated region, at least for *CIRBP* sequence.

### N. Putative protein members of the excising mechanism

To detect protein factors participating in the excising process, we performed RNA-pulldown assay using *CIRBP* RNA as a target. For immobilization of the RNA on streptavidin agarose, we attached a 4-fold repeat of streptavidin aptamer S1M (24) at the 3’-end of the sequence. To detect the RNA binding proteins that have an affinity to the RNA sequence, we performed 2D electrophoresis of the RNA-pulldown results using Coomassie Blue R250 staining (Figure 6). Major spots were then identified with MALDI-tof. As a control, we used RFP mRNA, also with a 4xS1M sequence attached at the 3’ end. According to our results, the list of proteins that interact with *CIRBP* mRNA consists of mitochondrial leucine-rich PPR-containing protein (LRPPRC), interleukin enhancer-binding factors 2 and 3 (ILF2, 3), heat shock 70 kDa proteins 1A/1B (HSPA1A or HSP70), G-rich sequence factor 1 (GRSF1) and heterogeneous nuclear ribonucleoprotein H1 (hnRNPH1).

**Fig. 6.**
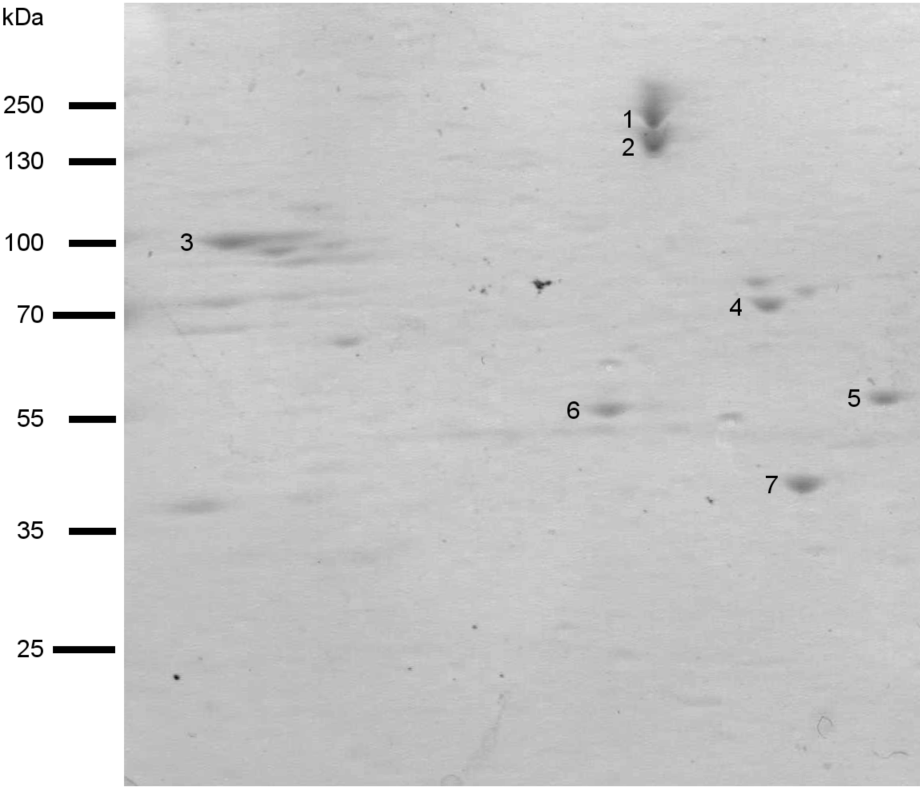
2D electrophoresis analysis of proteins associated with excising. The numbers denote individual spots. According to MALDI-tof, the numbered spots correspond to: 1, 2 - LRPPRC (glycated and non-glycated forms, correspondingly), 3 - ILF3, 4 - HSPA1A, 5 - GRSF1, 6 - hnRNPH1, 7 - ILF2.

Analysis of common pathways and interactions between the proteins revealed that HSPA1A, HSPA1B, ILF2, and ILF3 could specifically interact with each other (32, 33), which includes the formation of a multiprotein complex necessary for transcription at the beta-globin locus (34). ILF2 also can associate with hnRNPH1, while ILF3 can form a complex with LRPPRC (35). Finally, hnRNPH1 and LRPPRC could specifically recognize each other (36). Among these proteins, HSP70 can directly bind to U-rich RNA sequences (37, 38), which makes it a possible initiator of assembly of a multiprotein excising complex.

## Discussion

Eukaryotic cells have a vast repertoire of transcriptome and proteome diversity generating tools. Despite the apparent redundancy of this apparatus, we encountered a previously undescribed mechanism of mRNA post-transcriptional processing. The main aim of this study was to make sure that the observed phenomenon is genuine and not related to possible methodological or experimental artifacts.

The mechanism that we called excising manifested itself as a pattern of bands corresponding to different isoforms of *CIRBP* cDNA (see Figure 1). Interestingly, we discovered a similar appearance in *Mus musculus*, but with a different pattern (Supplementary Note 3 and Figure 7).

**Fig. 7.**
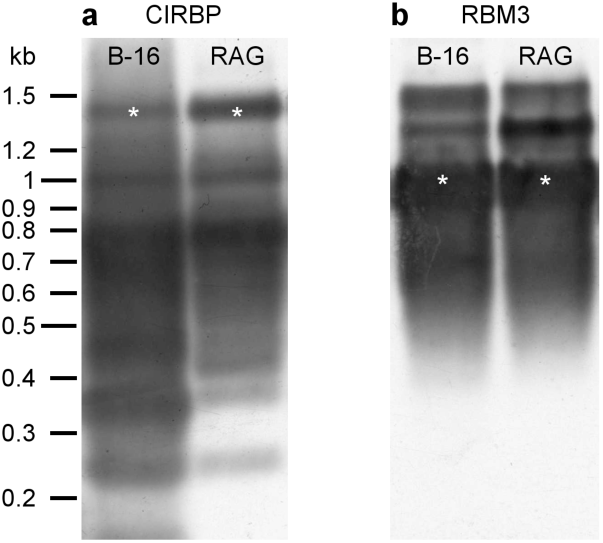
CIRBP and RBM3 transcripts diversity in *Mus Musculus*: southern blot analysis of RT-PCR products of CIRBP (**a**) and RBM3 (**b**) mRNA. Symbols “*” denote bands with expected lengths.

To prove that the observed transcript variety was not associated with the reverse transcription or PCR artifacts, we performed a northern blotting experiment. It showed the production of truncated mRNA variants from both the mature *CIRBP* transcript (vector RFC) and the same sequence lacking 3’-UTR (vector RFO, see Figure 3). These results confirmed that mRNA might be a direct target of the excising mechanism.

We have also created a construct to demonstrate that *CIRBP* excising leading to the deletion of a stop codon results in otherwise repressed *GFP* translation (see Figure 5e, h). Moreover, western blotting results verify the differential translation of constructs having and lacking 3’-UTR (see Figure 5a-c). It eliminates the possibility of artifacts associated with cloning protocol.

Another possible explanation for the observed phenomenon is the aberrant alternative splicing. We investigated this possibility using constructs containing the sequences of pre-mRNA (vector RFG) and mature transcript of *CIRBP* gene (vector RFC, see Figure 3a). In both cases, we observed similar band patterns (see Figure 3b), which supports the hypothesis that excising is a post-splicing event. We also discovered that spliceosome inhibitor isoginkgetin does not affect the patterns.

Earlier, a similar behavior was observed for *TGS101* and *FHIT* genes and was associated with aberrant alternative splicing via weak distant splicing sites (39, 40). Nevertheless, we discovered a large variety of aberrant *CIRBP* gene transcripts that differ from each other by 4-7 nucleotides. There were no clear consensus motifs at the junctions between the removed and retained sites (see Figure 2). It might be a result of a general disruption of the splicing process in the cell lines used in the study. However, the phenomenon was observed only for *CIRBP* mRNA, but not for the other genes (Figure S1). All these facts together indicate that excising is most likely a splicing-independent process.

Finally, excising is not a part of mRNA decay pathways. It seems to be a process of alternative processing of a pool of initial sequences rather than a stepwise degradation of a primal transcript. Further, post-excising mRNAs could be translated. While we observed protein products of processed mRNAs only in synthetic systems using expression vectors, there is no reason to exclude the possibility in natural systems. Note that the functionality of the protein products of excising raises certain doubts since both the deletion of significant sections of the sequence and the translation of the 3’-UTR can occur here. One can only assume the biological role of this process. We may face some adjuvant way of either CIRBP activity inhibition or generation of structural and functional diversity of its protein products. Moreover, the abundance of mitochondrial protein LRPPRC according to RNA-pulldown results might indicate the link between excising and cell metabolism activity.

Excising has a set of unique features, which distinguishes it as a standalone mechanism. First, it is a low-accuracy process. While the observed variety of transcripts would have to have a continuous distribution of electrophoretic mobility, in reality, it was discrete (see Figures 1, 3b, 4b). We believe that it might be associated with specific recognition and deletion of U-rich stretches, which alters the sequence making it less vulnerable to another round of the excising mechanism.

Second, according to MALDI-tof results excising seems to have a unique multiprotein complex responsible for the phenomenon (see Figure 6). All the proteins from the complex were shown to form specific pairs, but there is no published data on their joint participation in any specific metabolic process. Nevertheless, the involvement of detected proteins in the excising process should be experimentally validated. Moreover, there is a possibility of additional members of the multiprotein complex, which we could not identify. We plan to investigate both issues in the following works.

Finally, excising is a selective process. From all the genes that we have studied, only *CIRBP* was a target of the mechanism. Even *RBM3*, closely related to *CIRBP* (65% nucleic sequence identity), showed no signs of excising (see Figure S1). The identification of other genes vulnerable to the process is also an intriguing theme of future research.

The second mechanism that we have discovered is a phenomenon of incising. It involves transcript elongation due to duplication of regions of the sequence, in some cases accompanied by excising (see Figure 4b, c). Although we detected the process on the shortest construct studied (R3), other expression vectors were also the subjects of incising. We believe that this process is associated with the formation of stable secondary structures interacting with RNA-binding proteins. For example, the R3 vector sequence might form a pseudoknot structure with a length of 15 bp containing two consecutive G-U pairs (Figure 4f). According to Su et al., G-T mismatches may be a preferred site for protein side-chain intercalation (41). This hypothesis concurs with the fact that the pseudoknot-forming sequence was a part of the duplicated region (see Figure 4c, f and Supplementary Note 2).

## Conclusions

In this article, we present a previously undescribed mechanism of *CIRBP* mRNA post-transcriptional processing resulting in a remarkable but discrete variety of resulting transcripts. The process named by us “excising” differs from other mechanisms of transcriptome and proteome diversity generation. First, it seems to be associated with a unique multiprotein complex. Second, it is a low-accuracy process, where individual transcripts might differ from each other by 4-7 nucleotides. Finally, excising is a selective process, targeting genes according to not yet known criteria. The main observable effect of excising is the deletion of sequence regions of variable length from the transcripts, which could lead, among other possibilities, to 3’-UTR translation in case of stop-codon removal. However, in contrast to splicing, there were no consensus motifs at the junctions between the removed and retained sites. Several other features, such as immunity to splicing inhibitor or lack of intronic sequences effect, distinguish excising as a post-splicing and splicing-independent process.

## ACKNOWLEDGEMENTS

The work was supported by Russian Foundation for Basic Research, research project [18-29-01012]. The authors thank Rustam Ziganshin (Shemyakin and Ovchinnikov Institute of Bioorganic Chemistry) for proteomic research and valuable recommendations. The authors are grateful to the Member of Russian Academy of Sciences, Professor Pavel G. Georgiev for valuable discussion of this work.

## O. Supplementary Information

### Supplementary Note 1: Sequences of CIRBP transcript variants and RNA-Seq reads used for Figure 2

#### Series 500 clones 17, 35a, 35b, 38

ATGGCATCAGATGAAGGCAAACTTTTTGTTGGAGG GCTGAGTTTTGACACCAATGAGCAGTCGCTGGAGC AGGTCTTCTCAAAGTACGGACAGATCTCTGAAGTG GTGGTTGTGAAAGACAGGGAGACCCAGAGATCTCG GGGATTTGGGTTTGTCACCTTTGAGAACATTGACG ACGCTAAGGATGCCATGATGGCCATGAATGGGAAG TCTGTAGATGGACGGCAGATCCGAGTAGACCAGGC AGGCAAGTCGTCAGACAACCGATCCCGTGGGTACC GTGGTGGCTCTGCCGGGGGCCGGGGCTTCTTCCGT GGGGGCCGAGGACGGGGCCGTGGGTTCTCTAGAG GAGGAGGGGACCGAGGCTATGGGGGGAACCGGTT CGAGTCCATCTGTGCTGTGCGCCCCACAGTAGACG TGCAGACGTCCCTGAGAGGTTCTTGAAGATGTTTA TTTATATTGTCCTTTTTTACTGGAAGACGTACGCAT A

#### Series 500 clone 25

ATGGCATCAGATGAAGGCAAACTTTTTGTTGGAGG GCTGAGTTTTGACACCAATGAGCAGTCGCTGGAGC AGGTCTTCTCAAAGTACGGACAGATCTCTGAAGTG GTGGTTGTGAAAGACAGGGAGACCCAGAGATCTCG GGGATTTGGGTTTGTCACCTTTGAGAACATTGACG ACGCTAAGGATGCCATGATGGCCATGAATGGGAAG TCTGTAGATGGACGGCAGATCCGAGTAGACCAGGC AGGCAAGTCGTCAGACAACCGATCCCGTGGGTACC GTGGTGGCTCTGCCGGGGGCCGGGGCTTCTTCCGT GGGGGCCGAGGACGGGGCCGTGGGTTCTCTAGAG GAGGAGGGGACCGAGGCTATGGGGGGAACCGGTT CCAGCCCCTGCGTCCATCTGTGCTGTGCGCCCCACA GTAGACGTGCAGACGTCCCTGAGAGGTTCTTGAAG ATGTTTATTTATATTGTCCTTTTTTACTGGAAGACG TACGCATA

#### Series 500 clone 33

ATGGCATCAGATGAAGGCAAACTTTTTGTTGGAGG GCTGAGTTTTGACACCAATGAGCAGTCGCTGGAGC AGGTCTTCTCAAAGTACGGACAGATCTCTGAAGTG GTGGTTGTGAAAGACAGGGAGACCCAGAGATCTCG GGGATTTGGGTTTGTCACCTTTGAGAACATTGACG ACGCTAAGGATGCCATGATGGCCATGAATGGGAAG TCTGTAGATGGACGGCAGATCCGAGTAGACCAGGC AGGCAAGTCGTCAGACAACCGATCCCGTGGGTACC GTGGTGGCTCTGCCGGGGGCCGGGGCTTCTTCCGT GGGGGCCGAGGACGGGGCCGTGGGTTCTCTAGAG GAGGAGGGGACCGAGGCTATGGGGGGAACCGGTT CGAGCCCCTGCGTCCATCTGTGCTGTGCGCCCCAC AGTAGACGTGCAGACGTCCCTGAGAGGTTCTTGAA GATGTTTATTTATATTGTCCTTTTTTACTGGAAGAC GTACGCATA

#### Series 500 clone 34

ATGGCATCAGATGAAGGCAAACTTTTTGTTGGAGG GCTGAGTTTTGACACCAATGAGCAGTCGCTGGAGC AGGCCTTCTCAAAGTACGGACAGATCTCTGAAGTG GTGGTTGTGAAAGACAGGGAGACCCAGAGATCTCG GGGATTTGGGTTTGTCACCTTTGAGAACATTGACG ACGCTAAGGATGCCATGATGGCCATGAATGGGAAG TCTGTAGATGGACGGCAGATCCGAGTAGACCAGGC AGGCAAGTCGTCAGACAACCGATCCCGTGGGTACC GTGGTGGCTCTGCCGGGGGCCGGGGCTTCTTCCGT GGGGGCCGAGGACGGGGCCGTGGGTTCTCTAGAG GAGGAGGGGACCGAGGCTATGGGGGGAACCGGTT CGAGTCCTGCGTCCATCTGTGCTGTGCGCCCCACA GTAGACGTGCAGACGTCCCTGAGAGGTTCTTGAAG ATGTTTATTTATATTGCCCTTTTTTTACTGGAAGAC GTACGCATA

#### Series 500 clone 39

ATGGCATCAGATGAAGGCAAACTTTTTGTTGGAGGGCTGAGTTTTGACACCAATGAGCAGTCGCTGGAGCAGGTCTTCTCAAAGTACGGACAGATCTCTGAAGTGGTGGTTGTGAAAGACAGGGAGACCCAGAGATCTCGGGGATTTGGGTTTGTCACCTTTGAGAACATTGACGACGCTAAGGATGCCATGATGGCCATGAATGGGAAGTCTGTAGATGGACGGCAGATCCGAGTAGACCAGGCAGGCAAGTCGTCAGACAACCGATCCCGTGGGTACCGTGGTGGCTCTGCCGGGGGCCGGGGCTTCTTCCGTGGGGGCCGAGGACGGGGCCGTGGGTTCTCTAGAGGAGGAGGGGACCGAGGCTATGGGGGGAACCGGTTCGAGTCCAGGAGTGGGGGCTACCCAGCCCCTGCGTCCATCTGTGCTGTGCGCCCCACAGTAGACGTGCAGACGTCCCTGAGAGGTTCTTGAAGATGTTTATTTATATTGTCCTTTTTTACTGGAAGACGTACGCATA

#### Series 500 clone 42

ATGGCATCAGATGAAGGCAAACTTTTTGTTGGAGGGCTGAGTTTTGACACCAATGAGCAGTCGCTGGAGCAGGTCTTCTCAAAGTACGGACAGATCTCTGAAGTGGTGGTTGTGAAAGACAGGGAGACCCAGAGATCTCGGGGATTTGGGTTTGTCACCTTTGAGAACATTGACGACGCTAAGGATGCCATGATGGCCATGAATGGGAAGTCTGTAGATGGACGGCAGATCCGAGTAGACCAGGCAGGCAAGTCGTCAGACAACCGATCCCGTGGGTACCGTGGTGGCTCTGCCGGGGGCCGGGGCTTCTTCCGTGGGGGCCGAGGACGGGGCCGTGGGTTCTCTAGAGGAGGAGGGGACCGAGGCTATGGGGGGAACCGGTTCGAGTCCAGCCCCTGCGTCCATCTGTGCTGTGCGCCCCACAGTAGACGTGCAGACGTCCCTGAGAGGTTCTTGAAGATGTTTATTTATATTGTCCTTTTTTACTGGAAGACGTACGCATA

#### Series 500 clone 47

ATGGCATCAGATGAAGGCAAACTTTTTGTTGGAGG GCTGAGTTTTGACACCAATGAGCAGTCGCTGGAGC AGGTCTTCTCAAAGTACGGACAGATCTCTGAAGTG GTGGTTGTGAAAGACAGGGAGACCCAGAGATCTCG GGGATTTGGGTTTGTCACCTTTGAGAACATTGACG ACGCTAAGGATGCCATGATGGCCATGAATGGGAAG TCTGTAGATGGACGGCAGATCCGAGTAGACCAGGC AGGCAAGTCGTCAGACAACCGATCCCGTGGGTACC GTGGTGGCTCTGCCGGGGGCCGGGGCTTCTTCCGT GGGGGCCGAGGACGGGGCCGTGGGTTCTCTAGAG GAGGAGGGGACCGAGGCTATGGGGGGAACCGGTT CGAGTCCAGGAGTGGGGGCTACGGAGGCTCCAGA GACTACTATAGCAGCCCCTGCGTCCATCTGTGCTGT GCGCCCCACAGTAGACGTGCAGACGTCCCTGAGAG GTTCTTGAAGATGTTTATTTATATTGTCCTTTTTTAC TGGAAGACGTACGCATA

#### Series 1000 clone 14

ATGGCATCAGATGAAGGCAAACTTTTTGTTGGAGG GCTGAGTTTTGACACCAATGAGCAGTCGCTGGAGC GGGTCTTCTCAAAGTACGGACAGATCTCTGAAGTG GTGGTTGTGAAAGACAGGGAGACCCAGAGATCTCG GGGATTTGGGTTTGTCACCTTTGAGAACATTGACG ACGCTAAGGATGCCATGATGGCCATGAATGGGAAG TCTGTAGATGGACGGAGGCTCCAGAGACTACTATA GCAGCCGGAGTCAGAGTGGTGGCTACAGTGACCGG AGCTCGGGCGGGTCCTACAGAGACAGTTACGACAG TTACGCTACACACAACGAGTAAAAACCCTTCCTGC TCAAGATCGTCCTTCCAATGGCTGTGTGTTTAAAGA TTGTGGGAGCTTCGCTGAACGTTAATGTGTAGTAA ATGCACCTCCTTGTATTCCCACTTTCGTAGTCATTT CGGTTCTGATCTTGTCAAACCCAGCCTGACCGCTTC TGACGCCGGGATGGCCTCGTTACTAGACTTTTCTTT TTAAGGAAGTGCTGTTTTTTTTTTGAGGGTTTTCAA AACATTTTGAAAAGCATTTACTTTTTTGACCACGAG CCATGAGTTTTCAAAAAAATCGGGGGTTGTGTGGG TTTTTGGTTTTCGTTTTAGTTTTTGGTTGCGTTGCCT TTTTTTTTTTTTTAGTGGGGTTGGCCCCATGAAGTG GGTGCCCCACTCACTTCTCTGAGATCGAACGGACT GTGAATCCGCTCTTTGTCGGAAGCTGAGCAAGCTG TGGCTTTTTTCCAACTCCGTGTGACGTTTCTGAGTG TAGTGTGGTAGGACCCCGGCGGGTGTGGCAGCAAC TGCCCTGGAGCCCCAGCCCCTGCGTCCATCTGTGCT GTGCGCCCCACAGTAGACGTGCAGACGTCCCTGAG AGGTTCTTGAAGATGTTTATTTATATTGTCCTTTTTT ACTGGAAGACGTACGCATAGGATCCACCGGATCTA GATAATCGAATTCCCGCGGCCGCCATGGCGGCCGG GA

#### Series 1000 clone 20

ATGGCATCAGATGAAGGCAAACTTTTTGTTGGAGG GCTGAGTTTTGACACCAATGAGCAGTCGCTGGAGC AGGTCTTCTCAAAGTACGGACAGATCTCTGAAGTG GTGGTTGTGAAAGACAGGGAGACCCAGAGATCTCG GGGATTTGGGTTTGTCACCTTTGAGAACATTGACG ACGCTAAGGATGCCATGATGGCCATGAATGGGAAG TCTGTAGATGGACGGCAGATCCGAGTAGACCAGGC AGGCAAGCCGTCAGACAACCGATCCCGTGGGTACC GTGGTGGCTCTGCCGGGGGCCGGGGCTTCTTCCGT GGGGGCCGAGGACGGGGCCGTGGGTTCTCTAGAG GAGGAGGGGACCGAGGCTATGGGGGGAACCGGTT CGAGTCCAGGAGTGGGGGCTACGGAGGCTCCAGA GACTACTATAGCAGCCGGAGTCAGAGTGGTGGCTA CAGTGACCGGAGCTCGGGCGGGTCCTACAGAGACA GTTACGACAGTTACTAGACTTTTCTTTTTAAGGAAG TGCTGTTTTTTTTGAGGGTTTTCAAAACATTTTGAA AAGCATTTACTTTTTTGACCACGAGCCATGAGTTTT CAAAAAAATCGGGGGTTGTGTGGGTTTTTGGTTTTT GTTTTAGTTTTTGGTTGCGTTGCCTTTTTTTTTTTTA GTGGGGTTGGCCCCATGAAGTGGGTGCCCCACTCA CTTCTCTGAGATCGAACGGACTGTGAATCCGCTCTT TGTCGGAAGCTGAGCAAGCTGTGGCTTTTTTCCAA CTCCGTGTGACGTTTCTGAGTGTGGTGTGGTAGGA CCCCGGCGGGTGTGGCAGCAACTGCCCTGGAGCCC CAGCCCCTGCGTCCATCTGTGCTGTGCGCCCCACA GTAGACGTGCAGACGTCCCTGAGAGGTTCTTGAAG ATGTTTATTTATATTGTCCTTTTTTACTGGAAGACG TACGCATAGGATCCACCGGATCTAGATAATCGAAT TCCCGCGGCCGCCATGGCGGCCGGGAGCATGCGA

#### Series 1000 clone 24

ATGGCATCAGATGAAGGCAAACTTTTTGTTGGAGGGCTGAGTTTTGACACCAATGAGCAGTCGCTGGAGCAGGTCTTCTCAAAGTACGGACAGATCTCTGAAGTGGTGGTTGTGAAAGACAGGGAGACCCAGAGATCTCGGGGATTTGGGTTTGTCACCTTTGAGAACATTGACGACGCTAAGGATGCCATGATGGCCATGAATGGGAAGTCTGTAGATGGACGGCAGATCCGAGTAGACCAGGCAGGCAAGTCGTCAGACAACCCGATCCCGTGGGTACCGTGGTGGCTCTGCCGGGGGCCGAGGACGGGGCCGAGGACAGTTACGACAGTTACGCTACACACAACGAGTAAAAACCCTTCCTGCTCAAGATCGTCCTTCCAATGGCTGTGTGTTTAAAGATTGTGGGAGCTTCGCTGAACGTTAATGTGTAGTAAATGCACCTCCTTGTATTCCCACTTTCGTAGTCATTTCGGTTCTGATCTTGTCAAACCCAGCCTGACCGCTTCTGACGCCGGGATGGCCTCGTTACTAGACTTTTCTTTTTAAGGAAGTGCTGTTTTTTTTTTGAGGGTTTTCAAAACATTTTGAAAAGCATTTACTTTTTTGACCACGAGCCATGAGTTTTCAAAAAAATCGGGGGTTGTGTGGGTTTTTGGTTTTTGTTTTAGTTTTTGGTTGCGTTGCCTTTTTTTTTTTTAGTGGGGTTGGCCCCATGAAGTGGGTGCCCCACTCACTTCTCTGAGATCGAACGGACTGTGAATCCGCTCTTTGTCGGAAGCTGAGCAAGCTGTGGCTTTTTTCCAACTCCGTGTGACGTTTCTGAGTGTAGTGTGGTAGGACCCCGGCGGGTGTGGCAGCAACTGCCCTGGAGCCCCAGCCCCTGCGTCCATCTGTGCTGTGCGCCCCACAGTAGACGTGCAGACGTCCCTGAGAGGTTCTTGAAGATGTTTATTTATATTGTCCTTTTTTACTGGAAGACGTACGCATAGGATCCACCGGATCTAGATAATCGAATTCCCGCGGCCGCCATGGCGGCCGGGAG

#### Series 1000 clone 26

ATGGCATCAGATGAAGGCAAACTTTTTGTTGGAGG GCTGAGTTTTGACACCAATGAGCAGTCGCTGGAGC AGGTCTTCTCAAAGTACGGACAGATCTCTGAAGTG GTGGTTGTGAAAGACAGGGAGACCCAGAGATCTCG GGGATTTGGGTTTGTCACCTTTGAGAACATTGACG ACGCTAAGGATGCCATGATGGCCATGAATGGGAAG TCTGTAGATGGACGGCAGATCCGAGTAGACCAGGC AGGCAAGTCGTCAGACAACCGATCCCGTGGGTACC GTGGTGGCTCTGCCGGGGGCCGGGGCTTCTTCCGT GGGGGGACCGAGGCTATGGGGGGGGAACCGGTTC GAGTCCAGGAGTGGGGGGCTACGGAGGCTCCAGA GACTACTATAGCGGCCGGAGTCAGAGTGGTGGCTA CAGTGACCGGAGCTCGGGCGGGTCCTACAGAGACA GTTACGACAGTTACGCTACACACAACGAGTAAAAA CCCTTCCTGCTCAAGATCGTCCTTCCAATGGCTGTG TGTTTAAAGATTGTGGGAGCTTCGCTGAACGTTAA TGTGTAGTAAATGCACCTCCTTGTATTCCCACTTTC GTAGTCATTTCGGTTCTGATCTTGTCAAACCCAGCC TGACCGCTTCTGACGCCGGGATGGCCTCGTTACTA GACTTTTCTTTTTAAGGAAGTGCTGTTTTTTTTTTGA GGGTTTTCAAAACATTTTGAAAAGCATTTACTTTTT TGACCACGAGCCATGAGTTTTCAAAAAAATCGGGG GTTGTGTGGGTTTTTGGTTTTTGTTTTAGTTTTTGGT TGCGTTGCCTTTTTTTTTTTTAGTGGGGTTGGCCCC ATGAAGTGGGTGCCCCACTCACTTCTCTGAGATCG AACGGACTGTGAATCCGCTCTTTGTCGGAAGCTGG AGCCCCAGCCCCTGCGTCCATCTGTGCTGTGCGCCC CACAGTAGACGTGCAGACGTCCCTGAGAGGTTCTT GAAGATGTTTATTTATATTGTCCTTTTTTACTGGAA GACGTACGCATAGGATCCACCGGATCTAGATAATC ACTAGTGAATTCGCGGCCGCCTGCAGGTCGACCAT ATGGGAGAGCTCCCAACGCGTTGGATGCATAGCTT GAGTATTCTATAGTGTCACCTA

#### Series 1000 clone 27

ATGGCATCAGATGAAGGCAAACTTTTTGTTGGAGG GCTGAGTTTTGACACCAATGAGCAGTCGCTGGAGC AGGTCTTCTCAAAGTACGGACAGATCTCTGAAGTG GTGGTTGTGAAAGACAGGGAGACCCAGAGATCTCG GGGATTTGGGTTTGTCACCTTTGAGAACGTTGACG ACGCTAAGGATGCCATGATGGCCATGAATGGGAAG TCTGTAGATGGACGGCAGATCCGAGTAGACCAGGC AGGCAAGTCGTCAGACAACCGATCCCGTGGGTACC GTGGTGGCTCTGCCGGGGGCCGGGGCTTCTTCCGT GGGGGGCCGAGGACGGGGCCGTGGGTTCTCTAGA GGAGGAGGGGACCGAGGCTATGGGGGGAACCGGT TCGAGTCCAGGAGTGGGGGCTACGGAGGCTCCAGA GACTACTATAGCAGCCGGAGTCAGAGTGGTGGCTA CAGTGACCGGAGCTCGGGCGGGTCCTACAGAGACA GTTACGACAGTTACGCTACACACAACGAGTAAAAA CCCTTCCTGCTCAAGATCGTCCTTCCAATGGCTGTG TGTTTAAAGATTGTGGGAGCTTCGCTGTTTTTTTTT TTGAGGGTTTTCAAAACATTTTGAAAAGCATTTACT TTTTTGACCACGAGCCATGAGTTTTCAAAAAAATC GGGGGTTGTGTGGGTTTTTGGTTTTTGTTTTAGTTTT TGGTTGCGTTGCCTTTTTTTTTTTTTAGTGGGGTTGG CCCCATGAAGTGGGTGCCCCACTCACTTCTCTGAG ATCGAACGGACTGTGAATCCGCTCTTTGTCGGAAG CTGAGCAAGCTGTGGCTTTTTTTCCAACTCCGTGTG ACGTTTCTGAGTGTAGTGTGGTAGGACCCCGGCGG GTGTGGCAGCAACTGCCCTGGAGCCCCAGCCCCTG CGTCCATCTGTGCTGTGCGCCCCACAGTAGACGTG CAGACGTCCCTGAGAGGTTCTTGAAGATGTTTATTT ATATTGTCCTTTTTTACTGGAAGACGTACGCATAGG ATCCACCGGATCTAGATAATCGAATTCCCGCGGCC GCCATGGCGGCCGGGAGCATGCGACGTCGGGCCCA ATCGCCCCTTTTAGTAAGTGAA

#### Series 1000 clone 29

ATGGCATCAGATGAAGGCAAACTTTTTGTTGGAGG GCTGAGTTTTGACACCAATGAGCAGTCGCTGGAGC AGGTCTTCTCAAAGTACGGACAGATCTCTGAAGTG GTGGTTGTGAAAGACAGGGAGACCCAGAGATCTCG GGGATTTGGGTTTGTCACCTTTGAGAACATTGACG ACGCTAAGGATGCCATGATGGCCATGAATGGGAAG TCTGTAGATGGACGGCAGATCCGAGTAGACCAGGC AGGCAAGTCGTCAGACAACCGATCCCGTGGGTACC GTGGTGGCTCTGCCGGGGGGCCGGGGGCTTCTTCC GTGGGGGCCGAGGACGGGGCCGTGGGTTCTCTAGA GGAGGAGGGGACCGAGGCTATGGGGGGAACCGGT TCGAGTCCAGGAGTGGGGGCTACGGAGGCTCCAGA GACTACTATAGCAGCCGGAGTCAGAGTGGTGGCTA CAGTGACCGGAGCTCGGGCGGGTCCTACAGAGACA GTTACGACAGTTACTAGACTTTTCTTTTTAAGGAAG TGCTGTTTTTTTTTTTGAGGGTTTTCAAAACATTTTG AAAAGCATTTACTTTTTTGACCACGAGCCATGAGTT TTCAAAAAAATCGGGGGTTGTGTGGGTTTTTGGTTT TTGTTTTAGTTTTTGGTTGCGTTGCCTTTTTTTTTTT AGTGGGGTTGGCCCCATGAAGTGGGTGCCCCACTC ACTTCTCTGAGATCGAACGGACTGTGAATCCGCTC TTTGTCGGAAGCTGAGCAAGCTGTGGCTTTTTTCCA ACTCCGTGTGACGTTTCTGAGTGTAGTGTGGTAGG ACCCCGGCGGGTGTGGCAGCAACTGCCCTGGAGCC CCAGCCCCTGCGTCCATCTGTGCTGTGCGCCCCACA GTAGACGTGCAGACGTCCCTGAGAGGTTCTTGAAG ATGTTTATTTATATTGTCCTTTTTTACTGGAAGACG TACGCATAGGATCCACCGGATCTAGATAATCACTA GTGAATTCGCGGCCGCCTGCAGGTCGACCATATGG GAGAGCTCCCAACGCGTTGGATGCATAGCTTGAGT ATTCTATAGTGTCACCTAA

#### Series 1000 clone 31

ATGGCATCAGATGAAGGCAAACTTTTTGTTGGAGGGCTGAGTTTTGACACCAATGAGCAGTCGCTGGAGCAGGTCTTCTCAAAGTACGGACAGATCTCTGAAGTGGTGGTTGTGAAAGACAGGGAGACCCAGAGATCTCGGGGATTTGGGTTTGTCACCTTTGAGAACATTGACGACGCTAAGGATGCCATGATGGCCATGAATGGGAAGTCTGTAGATGGACGGCAGATCCGAGTAGACCAGGCAGGCAAGTAGCAGCCGGAGTCAGAGTGGTGGCTACAGTGACCGGAGCTCGGGCGGGTCCTACAGAGACAGTTACGACAGTTACGCTACACACAACGAGTAAAAACCCTTCCTGCTCAAGATCGTCCTTCCAATGGCTGTGTGTTTAAAGATTGTGGGAGCTTCGCTGAACGTTAATGTGTAGTAAATGCACCTCCTTGTATTCCCACTTTCGTAGTCATTTCGGTTCTGATCTTGTCAAACCCAGCCTGACCGCTTCTGACGCCGGGATGGCCTCGTTACTAGACTTTTCTTTTTAAGGAAGTGCTGTTTTTTTTTTGAGGGTTTTCAAAACATTTTGAAAAGCATTTACTTTTTTGACCACGAGCCATGAGTTTTCAAAAAAATCGGGGGTTGTGTGGGTTTTTGGTTTTTGTTTCAGTTTTTGGTTGCGTTGCCTTTTTTTTTTTAGTGGGGTTGGCCCCATGAAGTGGGTGCCCCACTCACTTCTCTGAGATCGAACGGACTGTGAATCCGCTCTTTGTCGGAAGCTGAGCAAGCTGTGGCTTTTTTCCAACTCCGTGTGACGTTTCTGAGTGTAGTGTGGTAGGACCCCGGCGGGTGTGGCAGCAACTGCCCTGGAGCCCCAGCCCCTGCGTCCATCTGTGCTGTGCGCCCCACAGTAGACGTGCAGACGTCCCTGAGAGGTTCTTGAAGATGTTCATTTATATTGTCCTTTTTTACTGGAAGACGTACGCATAGGATCCACCGGATCTAGATAATCGAATTCCCGCGGCCGCCATGGCGGCCGGGAG

#### Series 1000 clone 33

ATGGCATCAGATGAAGGCAAACTTTTTGTTGGAGG GCTGAGTTTTGACACCAATGAGCAGTCGCTGGAGC AGGTCTTCTCAAAGTACGGACAGATCTCTGAAGTG GTGGTTGTGAAAGACAGGGAGACCCAGAGATCTCG GGGATTTGGGTTTGTCACCTTTGAGAACATTGACG ACGCTAAGGATGCCATGATGGCCATGAATGGGAAG TCTGTAGATGGACGGCAGATCCGAGTAGACCAGGC AGGCAAGTCGTCAGACAACCGATCCCGTGGGTACC GTGGTGGCTCTGCCGGGGGCCGGGGCTTCTTCCGT GGGGGCCGAGGACGGGGGCCGTGGGTTCTCTAGA GGAGGAGGGGACCGAGGCTATGGGGGGAACCGGT TTCGAGTCCAGGAGTGGGGGCTACGGAGGCTCCAG AGACTACTATAGCAGCCGGAGTCAGAGTGGTGGCT ACAGTGACCGGAGCTCGGGCGGGTCCTACAGAGAC AGTTACGACAGTTACGCTACACACAACGAGTAAAA ACCCTTCCTGCTCAAGATCGTCCTTCCAATGGCTGT GTGTTTAAAGATTGTGGGAGCTTCGCTGAACGTTA ATGTGTAGTAAATGCACCTCCTCGTTACTAGACTTT TCTTTTTAAGGAAGTGCTGTTTTTTTTTTGAGGGTTT TCAAAACATTTTGAAAAGCATTTACTTTTTTGACCA CGAGCCATGAGTTTTCAAAAAAATCGGGGGTTGTG TGGGTTTTTGGTTTTTGTTTTAGTTTTTGGTTGCGTT GCCTTTTTTTTTTTTAGTGGGGTTGGCCCCATGAAG TGGGTGCCCCACTCACTTCTCTGAGATCGAACGGA CTGTGAATCCGCTCTTTGTCGGAAGCTGAGCAAGC TGTGGCTTTTTTCCAACTCCGTGTGACGTTTCTGAG TGTAGTGTGGTAGGACCCCGGCGGGTGTGGCAGCA ACTGCCCTGGAGCCCCAGCCCCTGCGTCCATCTGT GCTGTGCGCCCCACAGTAGACGTGCAGACGTCCCT GAGAGGTTCTTGAAGATGTTTATTTATATTGTCCTT TTTTACTGGAAGACGTACGCATAGGATCCACCGGA TCTTTTCTCGAATTCCCGCGGCCGCCATGGCGGCCG GGAGCA

#### Series 1000 clone 48

ATGGCATCAGATGAAGGCAAACTTTTTGTTGGAGG GCTGAGTTTTGACACCAATGAGCAGTCGCTGGAGC AGGTCTTCTCAAAGTACGGACAGATCTCTGAAGTG GTGGTTGTGAAAGACAGGGAGACCCAGAGATCTCG GGGATTTGGGTTTGTCACCTTTGAGAACATTGACG ACGCTAAGGATGCCATGATGGCCATGAATGGGAAG TCTGTAGATGGACGGCAGATCCGAGTAGACCAGGC AGGCAAGTCGTCAGACAACCGATCCCGTGGGTACC GTGGTGGCTCTGCCGGAGGCTCCAGAGACTACTAT AGCAGCCGGAGTCAGAGTGGTGGCTACAGTGACCG GAGCTCGGGCGGGTCCTACAGAGACAGTTACGACA GTTACGCTACACACAACGAGTAAAACCCTTCCTGC TCAAGATCGTCCTTCCAATGGCTGTGTGTTTAAAGA TTGTGGGAGCTTCGCTGAACGTTAATGTGTAGTAA ATGCACCTCCTTGTATTCCCACTTTCGTAGTCATTT CGGTTCTGATCTTGTCAAACCCAGCCTGACCGCTTC TGACGCCGGGATGGCCTCGTTACTAGACTTTTCTTT TTAAGGAAGTGCTGTTTTTTTTTTGAGGGTTTTCAA AACATTTTGAAAAGCATTTACTTTTTTGACCACGAG CCATGAGTTTTCAAAAAAATCGGGGGTTGTGTGGG TTTTTGGTTTTTGTTTTAGTTTTTGGTTGCGTTGCCT TTTTTTTTTTTAGTGGGGTTGGCCCCATGAAGTGGG TGCCCCACTCACTTCTCTGAGATCGAACGGACTGT GAATCCGCTCTTTGTCGGAAGCTGAGCAAGCTGTG GCTTTTTTCCAACTCCGTGTGACGTTTCTGAGTGTA GTGTGGTAGGACCCCGGCGGGTGTGGCAGCAGCTG CCCTGGAGCCCCAGCCCCTGCGTCCATCTGTGCTGT GCGCCCCACAGTAGACGTGCAGACGTCCCTGAGAG GTTCTTGAAGATGTTTATTTATATTGTCCTTTTTTAC TGGAAGACGTACGCATAGGATCCACCGGATCTAGA TAATCGAATTCCCGCGGCCGCCATGGCGGCCGGGA G

#### SRR1264355.20255217

GGAGGCTCGGGTCGTTGTGGCCGCCATGGCATCAGATGAAGGCAAACTTTTTGTTGGAGGGCTGAGTTTTGACACCAATGAGCAGTCGCTGGAGCAGGTCT

#### SRR1264355.53400006

GAGTTTTGACACCAATGAGCAGTCGCTGGAGCAGGTCTTCTCAAAGTACGGACAGATCTCGGGGATTTGGGTTTGTCACCTTTGAGAACATTGACGAGATC

#### SRR1264355.9124453

CATGATGGCCATGAATGGGAAGTCTGTAGATGGACGGCAGATCCGAGTAGACCAGGCAGGCAAGTCGTCAGACAACCGATCCCGAGGACGGGGCCGTGGGA

#### SRR1264355.43188086

GTCCTACAGAGACAGTTACGACAGTTACGTCTAGAAGCAGGCCAGAGAGCAGAGGCACGTGGCATCCCAGGGCGACCTCAGACGGCCAGCCGGTTAGCTAG

#### SRR1264355.47876842

GTTTAAAGATTGTGGGAGCTTCGCTGAACGTTAATGTGTAGTTGAAGGGGCCCGGCCAGGACTCGGGGAAGGGTGGCCTGAGAGCAGCGATGACCTCTGGG

#### SRR1264355.47876842/2

TAAAAACCCTTCCTGCTCAAGATCGTCCTTCCAATGGCTGTGTGTTTAAAGATTGTGGGAGCTTCGCTGAACGTTAATGTGTAGTTGAAGGGGCCCGGCCA

#### SRR1264355.1560728

ATCGTCCTTCCAATGGCTGTGTGTTTAAAGATTGTGGGAGCTTCGCTGAACGTTAATGTGCTGTTTTTTTTTGAGGGTTTTCAAAACATTTTGAAAAGCAT

### Supplementary Note 2: Sequences of transcript variants “∘” and “∼”

Gray: *GFP* sequence; gray and bold: duplicated segment of *GFP* sequence; black and bold: EcoRV restriction site; light-gray: *CIRBP* sequence; light-gray and bold: duplicated segment of *CIRBP* sequence.

#### Transcript variant “∘”

CGGGGCGGGGAGGCGAGGGCGACGCCGACTACGGCAAGCTGGAGATCAAGTTCATCTGCACCACCGGCAAGCTGCCCGTGCCCTGGCCCACCCTGGTGACCACCCTCTGCTACGGCATCCAGTGCTTCGCCCGCTACCCCGAGCACATGAAGATGAACGACTTCTTCAAGAGCGCCATGCCCGAGGGCTACATCCAGGAGCGCACCATCC AGTTCC **GATT** GCTGAACGTTAATGTGTAGTAAA TGCACCTCCTTGTATTCCCACTTTCGTAGTCATTTC GGTTCTGATCTTGTCAAACCCAG CCTGACCGCTT CTGACGCCGGGATGGCCTC **AATC** AGGACGA CGGCAAGTACAAGACCCGCGGCGAGGTGAAGTTCGAGGGCGACACCCTGGTGAACCGCATCGAGCTGGAGGGCAAGGACTTCAAGGAGGACGGCAACATCCTGGGCCACAAGCTGGAGTACAGCTTCAACAGCCACAACGTGTACATCCGCCCCGACAAGGCCA ACAACGGCCTGGAGGCTAACTTCAAGACCCGCCACAACATCGAGGGCGGCGGCGTGCAGCTGGCCGACCACTACCAGACCAACGTGCCCCTGGGCGAC GG CCCCGTGCTGGTCCCCATCAACCACTACCTGAGCACTCAGACCAAGATCAGCAAGGACCGCAACGAGGCCCGCGACCACATGGTGCTCCTGGAGTCCTTCAGCGCCTGCTGCCACACCCACGGCATGGACGAGCTGT ACAG CCTGACCGCTTCTGACGCCGGGATGGC CTC **AATC** AGGACGACGGCAAGTACAAGACCCGCGGCGAGGTGAAGTTCGAGGGCGACACCCTGTGAACCGCATCGAGCTGAAGGGCAAGGACTTCAAGGAGGACGGCAACATCCTGGCCACAAGCTGGAGTACAGCTTCAACAGCCACAACGTGTACATCCGCCCCGACAGGCCAACAACGGGCCTGAGGCTAACTTCAAGACCCGCCACACATCGAGGGCGGCGGCGTGCAGCTGGCCGACCACTACCAGACCAACGTGCCCTGGGCGACGG

#### Transcript variant “∼”

GGAGGCGAGGCGACGCCGACTACGGCAAGCTGGAGATCAAGTTCATCTGCACCACCGGCAAGCTGCCCGTGCCCTGGCCCACCCTGGTGACCACCCTCTGCTACGGCATCCAGTGCTTCGCCCGCTACCCCGAGCACATGAAGATGAACGACTTCTTCAAGAGCGCCATGCCCGAGGGCTACATCCA CCTGACCGCTTCTGACGCCGGGATGGCCTC **AATC** AGGACGACGGCAAGTACAAGACCCGCGGCGAGGTGAAGTTCGAGGGCGACACCCTGGTGAACCGCATCGAGCTGAAGGGCAAGGACTTCAAGGAGGACGGCAACATCCTGGGCCACAAGCTGGAGTACAGCTTCAACAGCCACAACGTGTACATCCGCCCCGACAAGGCCAACAACGGCCTGGAGGCTAACTTCAAGACCCGCCACAACATCGAGGGCGGCGGCGTGCAGCTGGCCGACCACTACCAGACCAACGTGCCCCTGGGCGACGGCCCCGTGCTGATCCCCATCAACCACTACCTGAGCACTCAGACCAAGATCAGCAAGGACCGCAACGAGGCCCGCGACCACATGGTGCTCCTGGAGTCCTTCAGCGCCTGCTGCCACACCCACGGCATGGACGAG CTG-TACAG CCTGACCGCTTCTGACGCCGGGATGGC CTC **AATC** AGGACGACGGCAAGTACAAGACCCGCGGCGAGGTGAAGTTCGAGGGCGACACCCTGGTGAACCGCATCGAGCTGAAGGGCAAGGACTTCAAGGAGGACGGCAACATCCTGGGCCACAAGCTGGAGTACAGCTTCAACAGCCACAACGTGTACATCCGCCCCGACAAGGCCAACAACGGCCTGTAGGCTAACTTCAAGACCCGCCACAACATCGAGGGCGGCGGCGTGCAGCTGGCCGACCACTACCAGACCAACGTGCCCCTGGGCGACGGCCCGTGCTGATCCCCATCAACCACTACCTGAGCACTCAGACCAGATCAGCAAGGACCGCAACGAGGCCCGCGACCACATGGTGCTCCCGGGAGTCCTTCAGCGCCTGCTGCCCCACCCACGGCATGTACGAG

### Supplementary Note 3: *CIRBP* and *RBM3* transcript variants in *Mus Musculus*

The total RNA was isolated from mouse cell lines B-16 (melanoma) and RAG (renal adenocarcinoma).

We used RT-PCR with two sets of primers, Cirb_M(f) vs. Cirb_exo7(r) and RbmMm(f) vs. RbmMm2(r), to study the diversity of CIRBP and RBM3 transcript variants, correspondingly. The sequences of the primers are listed in Table 1. The resulting PCR products were blotted to a nylon membrane, and southern hybridization with CIRBP-specific ZM1 probe and RBM3-specific ZM2 probe was performed. The probes exposed a variety of transcript variants with lengths less than expected for CIRBP gene (Figure 7a), but not for RBM3 (Figure 7b).

